# Computational analysis reveals the requirement for cell-cell interaction in liver repopulation by transplanted hepatocytes

**DOI:** 10.64898/2026.05.06.723162

**Authors:** Daiki Ikuta, Rei Tamaki, Sota Wada, Kotaro Onishi, Masaki Nishikawa, Yasuyuki Sakai, Takeshi Katsuda

## Abstract

Hepatocyte transplantation is a promising alternative to liver transplantation; however, it currently serves only as a temporary treatment until a donor organ becomes available. In contrast, animal studies have demonstrated “liver repopulation”, a phenomenon in which transplanted hepatocytes progressively replace host hepatocytes. Despite extensive documentation, the mechanisms driving this process remain poorly understood. More fundamentally, it remains unclear whether liver repopulation is driven by active cell–cell interactions between host and transplanted hepatocytes that induce host cell death, or whether it can be explained solely by intrinsic differences in proliferation and survival between these populations. To address this, we performed computational simulations using an agent-based model constrained by experimental data from repopulation in uninjured rat livers. Our analysis reveals that host hepatocyte death rate is the dominant determinant of repopulation kinetics, whereas variations in proliferation rate have only a limited impact. Notably, experimentally observed repopulation dynamics were only reproduced when cell– cell interactions were incorporated, or alternatively when host hepatocyte lifespan was set to unrealistically short values (approximately 25 days). These findings support a model in which cell– cell interactions play a critical role in efficient liver repopulation. More broadly, this study establishes a conceptual and computational framework for evaluating the requirement for cell–cell interactions in tissue replacement, even in the absence of a defined molecular mechanism.

## 1. Introduction

Hepatocyte transplantation has emerged as a promising alternative to liver transplantation, aiming to address donor shortage [1,2]. Its therapeutic basis lies in “liver repopulation,” a process in which transplanted hepatocytes replace host hepatocytes and restore liver function. Clinically, hepatocyte transplantation has shown efficacy in the treatment of genetic metabolic disorders and acute liver failure.

Despite this potential, hepatocyte transplantation is currently regarded as a temporary therapy, primarily serving as a bridge until donor organs become available. This limitation largely reflects the low efficiency of liver repopulation in humans, where transplanted hepatocytes typically replace less than 5% of liver mass [2]. Because restoration of liver function generally requires replacement of at least 10–30% of hepatocytes, depending on the disease context [3], this level of repopulation is often insufficient. In contrast, animal models frequently exhibit substantially higher repopulation efficiency, often exceeding 30% and leading to functional recovery [4]. This discrepancy underscores the need to better understand the mechanisms governing efficient liver repopulation.

Liver repopulation can be conceptualized as a two-step process: (1) engraftment of transplanted hepatocytes into the liver parenchyma and (2) subsequent replacement of host hepatocytes by engrafted cells [5]. The engraftment phase has been relatively well characterized and involves overcoming barriers such as sinusoidal endothelial traversal [6] and innate immune responses, both of which have been targeted to improve engraftment efficiency [6,7]. In contrast, the mechanisms underlying the replacement phase remain poorly understood.

A fundamental but unresolved question is whether hepatocyte replacement arises simply from the selective expansion of proliferative, healthy cells, or whether it is actively driven by cell–cell interactions between host and transplanted hepatocytes. In most experimental settings, repopulation is observed when healthy hepatocytes are introduced into injured livers, where host hepatocytes exhibit impaired proliferation and increased cell death. This observation is consistent with the intuitive hypothesis that transplanted hepatocytes outcompete host cells through a passive selective advantage [8].

Alternatively, several studies have suggested that active cell–cell interactions may contribute to hepatocyte replacement, including mechanisms related to cell competition [9,10], a process in which less fit cells are eliminated upon interaction with fitter neighbors [11–13]. For example, Oertel et al. reported increased apoptosis of host hepatocytes and enhanced proliferation of transplanted hepatocytes at the interface between the two populations [9]. However, this study was performed in a largely uninjured liver using fetal hepatocytes with high proliferative capacity, making it difficult to distinguish interaction-driven effects from simple expansion into space generated by physiological turnover.

Direct experimental evaluation of cell–cell interactions in this context remains challenging, largely due to the lack of a clear molecular framework. Even in the case of cell competition, a relatively well-documented phenomenon, key concepts such as “fitness” remain poorly defined, limiting the quantitative assessment of its contribution to liver repopulation.

As a complementary approach, computational modeling provides a means to test whether cell–cell interactions are required to explain observed repopulation dynamics. Agent-based models (ABMs) simulate the behavior of individual cells based on simple rules, enabling complex tissue-level behavior to emerge from local interactions [14,15]. In liver biology, ABMs have been applied to study metabolic regulation [16], regeneration after injury [17], and fibrosis progression [18,19]. Importantly, ABMs are well suited for systems with limited mechanistic information, as they allow direct evaluation of conceptual hypotheses.

In this study, we used an ABM-based framework to assess whether and to what extent cell– cell interactions are required for liver repopulation. We modeled cell–cell interaction as a local increase in the probability of apoptosis in host hepatocytes upon contact with transplanted cells. Using this simplified formulation, we show that experimentally observed repopulation kinetics cannot be reproduced without incorporating cell–cell interactions, supporting a critical role for such interactions in efficient liver repopulation.

## 2. Methods

### 2.1. Cell replacement model

A liver regeneration model was constructed using an agent-based modeling (ABM) framework. A schematic overview of the model is shown in **Figure 1A**. The computational domain consists of a 100 × 100 grid (10,000 sites), where each grid point can be occupied by a single cell.

**Figure 1.**
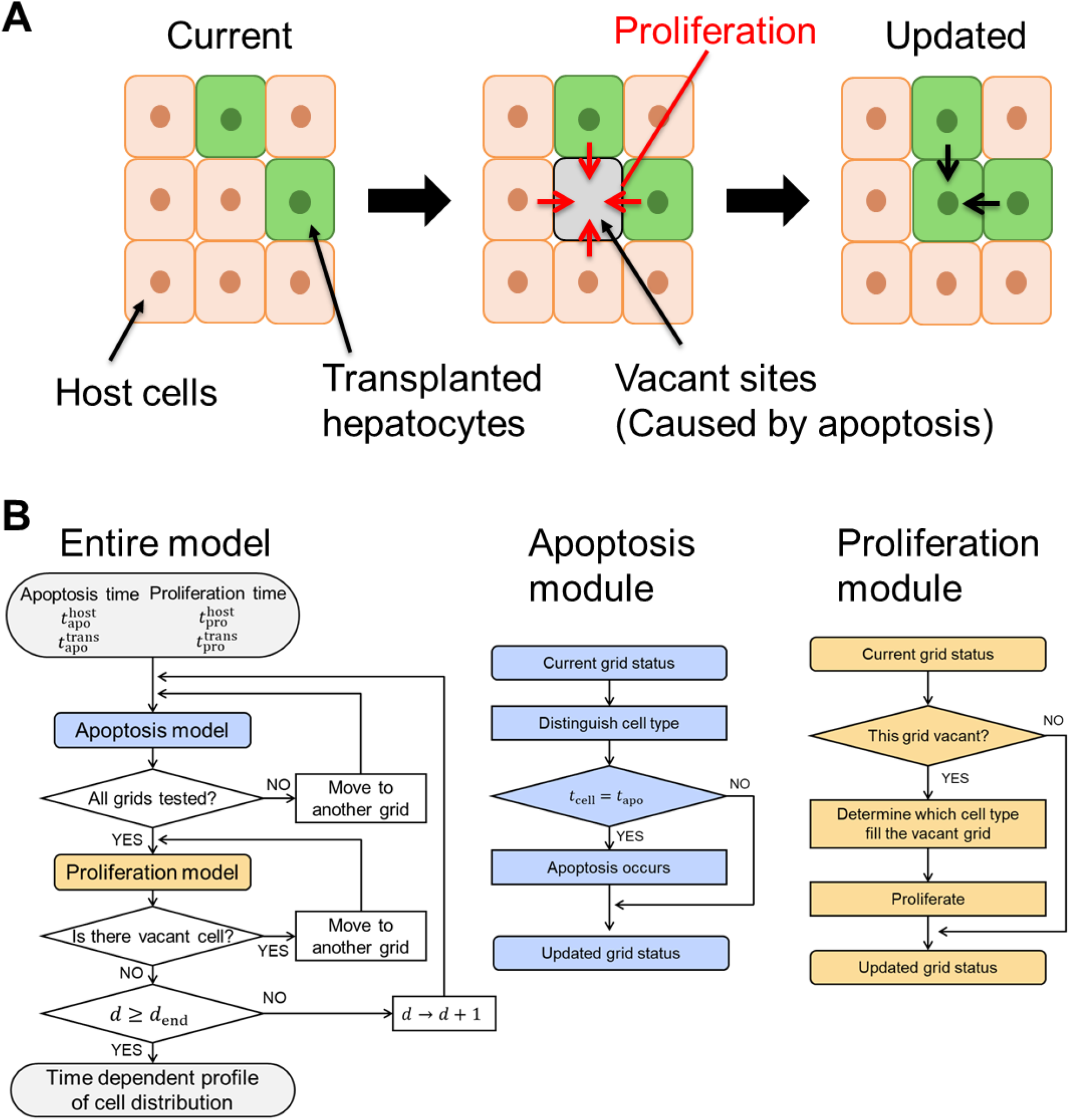
Overview of the cell replacement model. (A) Schematic illustration of the replacement process. Host hepatocytes (orange) and transplanted hepatocytes (green) occupy a discrete grid. Apoptosis generates vacant sites, which are subsequently filled by neighboring cells through space-dependent proliferation. (B) Flowchart of the simulation algorithm. At each time step, all grid sites are updated through apoptosis and proliferation modules, resulting in the time-dependent evolution of cell distribution.

Each cell is characterized by three state variables: the time elapsed since the last division, *t*_cell_ [h], the time to apoptosis after the last division, (i.e., cellular “lifespan”), *t*_apo_ [h], and a proliferation parameter, *t*_pro_ [h]. For host hepatocytes, the initial values of *t*_cell_ were assigned from a uniform distribution over the range [0, *t*_apo_).

The system evolves in discrete time steps of one day. At each simulated day *d* [d], all grid sites are updated through two processes: apoptosis and space-filling proliferation (**Figure 1B**). In the apoptosis module, any occupied grid in which *t*_cell_ = *t*_apo_ is cleared, representing cell death.

In the proliferation module, each vacant grid is filled by one of its neighboring cells (host or transplanted). The probability that a vacant grid is occupied by a host cell *p*^host^ [–] or a transplanted cell *p*^trans^ [–] is defined as:

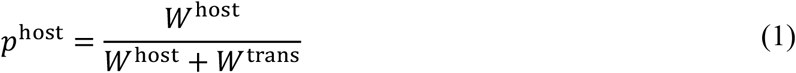

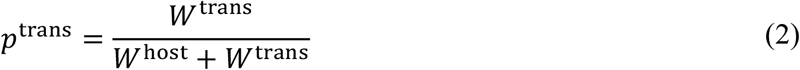

where the corresponding weights *W* reflect the relative proliferative capacity of neighboring cells. Specifically, the weights are defined as the sum of proliferation capacities *C*_pro_ of adjacent cells:

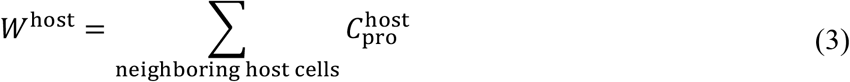

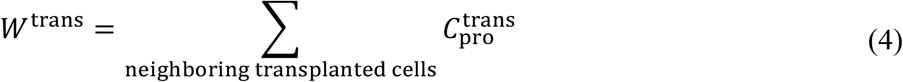

Here, the proliferation capacity is defined as the reciprocal of the proliferation parameter *C*_pro_ = 1⁄*t*_pro_.

Notably, proliferation is not explicitly modeled as a time-resolved cell division process. Instead, the proliferation parameter *t*_pro_ defines a relative growth advantage that determines the probability of occupying vacant space.

After applying the apoptosis module to all occupied grids and the proliferation module to all vacant grids, the simulation advances to the next day (*d* → *d* + 1). This process is repeated until the simulation reaches the predefined endpoint *d*_end_, yielding the time-dependent evolution of cell distribution.

### 2.2. Cell–cell interaction model

Cell–cell interaction was incorporated into the model as a local increase in the probability of apoptosis of host hepatocytes upon contact with transplanted cells (**Figure 2**). Specifically, on each simulated day, host cells that are in direct contact with transplanted cells undergo apoptosis with a probability *p*_apo_ [–], resulting in the corresponding grid becoming vacant.

**Figure 2.**
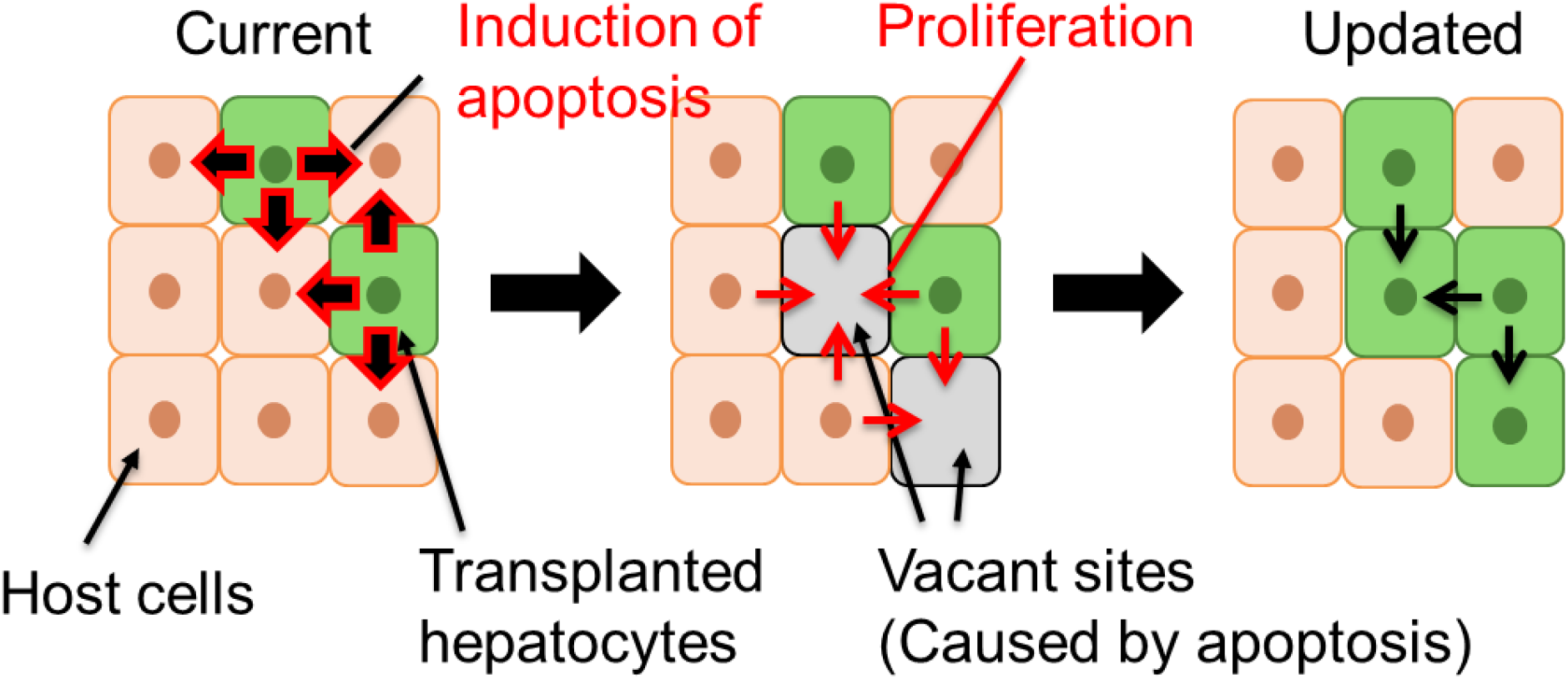
Schematic of the cell-cell interaction model. Transplanted hepatocytes induce apoptosis of adjacent host cells, generating additional vacant sites at the interface between the two populations. These sites are subsequently filled by neighboring cells, leading to enhanced expansion of transplanted hepatocytes.

The probability of apoptosis is defined as:

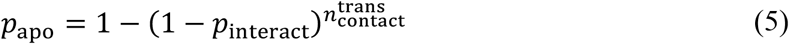

where *p*_interact_ [–] represents the per-contact probability that interaction with a transplanted cell induces apoptosis, and 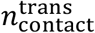 denotes the number of neighboring transplanted cells in contact with the host cell.

This formulation assumes that each contact independently contributes to apoptosis induction, such that the overall probability increases with the number of interacting transplanted cells. Thus, the model captures a local, contact-dependent effect of transplanted cells on host cell survival. This assumption is motivated by previous observations that host cell apoptosis is enriched at the interface between host and transplanted hepatocytes [9], although the underlying molecular mechanisms remain undefined.

### 2.3. Parameters and data set

Simulation results were compared with experimental data on hepatocyte transplantation and liver repopulation reported by Oertel et al., who examined the kinetics of liver repopulation following transplantation of embryonic day (ED) 14 fetal liver cells into adult rat livers [9]. Because the host liver in this study is uninjured, we assumed a homeostatic state in which hepatocyte turnover is balanced. Based on previous estimates of hepatocyte turnover time in adult rodent livers (200–400 days) [20–22], we set both the apoptosis time and proliferation parameter of host hepatocytes to 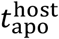 and 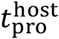 to 7200 h.

In contrast, fetal hepatocytes exhibit rapid growth. Therefore, apoptosis of transplanted cells was neglected 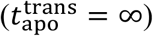. The proliferation parameter of transplanted hepatocytes was estimated from the volumetric growth rate of mouse fetal liver reported by Ogoke et al. [23], yielding 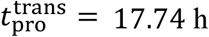. Notably, these parameter choices are intentionally biased in favor of transplanted cells, providing a conservative test of whether repopulation can be explained without cell–cell interactions.

The initial number of transplanted cells, 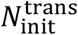, was defined as:

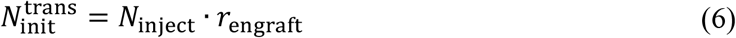

where *N*_inject_ represents the total number of administered cells and *r*_engraft_ denotes the engraftment ratio. In the study by Oertel et al., 4 − 6 × 10^7^ unfractionated fetal liver cells were transplanted, of which approximately 2% were hepatocyte progenitors. Accordingly, the number of cells with repopulation capacity was estimated to be *N*_inject_ ≅ 1 × 10^6^ cells per animal. Estimated values of *r*_engraft_ vary across studies [8,9,24,25], typically ranging from 0-10%, with maximum values up to 30% [24]. Therefore, we considered a range of *r*_engraft_ ∈ [0, 0.3].

Assuming that an adult rat liver contains approximately 1 × 10^9^ hepatocytes, the maximum initial fraction of transplanted cells corresponds to ∼0.03% (i.e. 3 × 10^5^ cells). To reflect this condition, we used a 100 × 100 grid (10,000 sites) and initialized the system with three transplanted cells (0.03%), placed at random positions. The repopulation ratio, *r*_repop_ [%] was defined as:

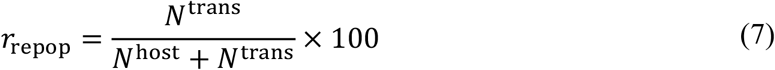

where *N*^trans^ and *N*^host^ [–] denote the numbers of transplanted and host cells, respectively.

All simulations were repeated 100 times with randomly assigned initial positions of transplanted cells, and the mean and standard deviation were calculated.

### 2.4. Sensitivity analysis

A sensitivity analysis was performed to assess the influence of key model parameters on the repopulation outcome. Specifically, the host cell properties 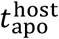 and 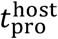 as well as the engraftment ratio *r*_engraft_, were systematically varied. For each parameter set, simulations were repeated 20 times with random initial conditions, and the average repopulation ratio at 180 days was calculated.

## 3. Results

Using an ABM approach, we developed a spatially explicit and dynamic model to describe liver repopulation by transplanted hepatocytes (**Figure 3A**). In this model, cell apoptosis is governed by cellular lifespan. The basal lifespan of host hepatocytes 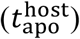 was set to 300 days, consistent with the estimated turnover time of adult rodent hepatocytes under homeostatic conditions [20–22].

**Figure 3.**
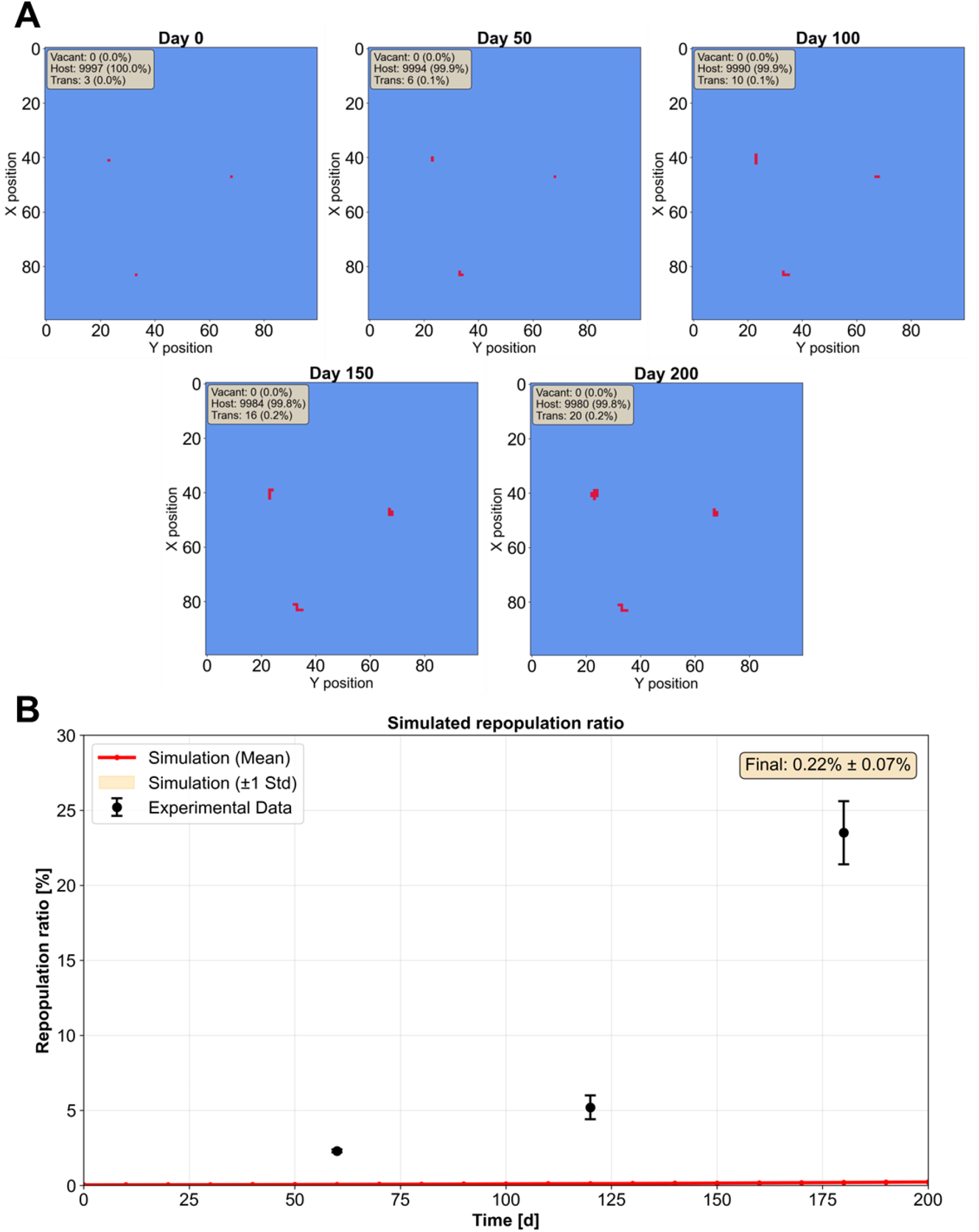
Simulation results under basal parameter settings 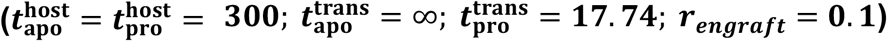. (A) Representative snapshots of simulated repopulation at indicated time points. Red and blue grids represent transplanted and host hepatocytes, respectively. (B) Temporal evolution of repopulation ratio. Simulation results (mean ± SD, n = 100 repeats) are shown together with experimental data.

### 3.1. Cell-cell interaction is necessary for repopulation in an uninjured host liver

We assumed that the driving force of repopulation arises from differences between host and transplanted cells in proliferative capacity (*C*_pro_ = 1⁄*t*_pro_) and apoptosis rate (represented by *t*_apo_). In this simulation, transplanted fetal hepatocytes were assigned a substantially higher proliferative capacity, with a doubling time 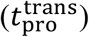 of 17.74 h, based on reported kinetics of mouse fetal liver growth [23]. Because apoptosis rates for fetal hepatocytes are not well defined in the literature, this parameter was left unconstrained.

To establish a null model lacking cell–cell interaction, we set 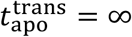, thereby maximizing the survival advantage of transplanted cells. Under this condition, host cell death occurs solely at the end of intrinsic lifespan. Notably, even under this highly favorable assumption, the simulated repopulation efficiency markedly deviated from experimental observations (**Figure 3B**), indicating that proliferative advantage alone is insufficient.

We next examined system behavior by varying 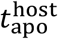 and 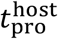. Because reducing 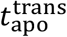 further decreases repopulation efficiency, this parameter was fixed at ∞. Sensitivity analysis revealed that the final replacement ratio is strongly influenced by host cell lifespan 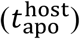, but is largely insensitive to host proliferative capacity 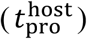 (**Figure 4A**). Temporal simulations, assuming an engraftment ratio of 0.1 (an optimistic estimate [9]), showed that 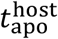 would need to be shorter than 10 days to reproduce experimentally observed repopulation kinetics (**Figure 4B**). However, hepatocyte turnover under homeostasis is typically estimated to be on the order of 200– 400 days [20–22]. Thus, the required condition 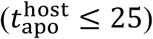 is physiologically unrealistic, further supporting the conclusion that proliferative advantage alone cannot account for repopulation dynamics.

**Figure 4.**
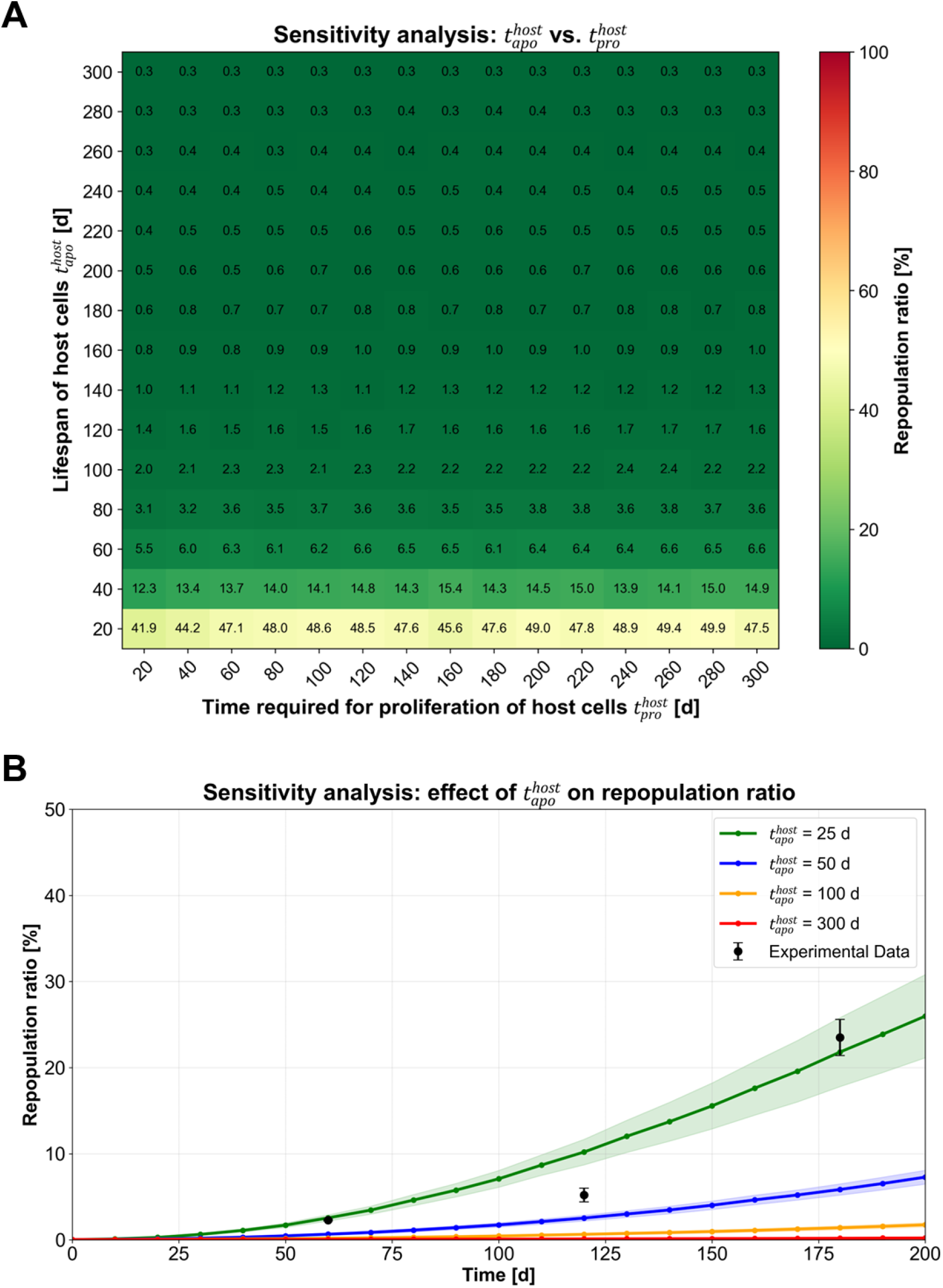
Sensitivity analysis of host cell properties. (A) Heatmap of repopulation ratio obtained by systematically varying 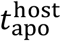 and 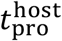. Each grid represents the simulated repopulation ratio. *r*_*engraft*_ was fixed to 0.1. (B) Temporal evolution of repopulation ratio for different values of 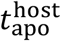 values.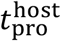 and *r*_*engraft*_ were fixed to 300 and 0.1, respectively.

We also evaluated the impact of engraftment efficiency (*r*_engraft_). Sensitivity analysis demonstrated that even under highly favorable conditions (*r*_engraft_ = 0.3), the model still requires 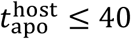 to match observed repopulation levels—again far shorter than physiological turnover rates (**Figure 5A**). Consistently, temporal simulations with 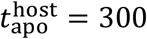 showed that even *r*_engraft_ = 1, an unrealistically favorable assumption, results in repopulation kinetics substantially slower than those observed experimentally (**Figure 5B**).

**Figure 5.**
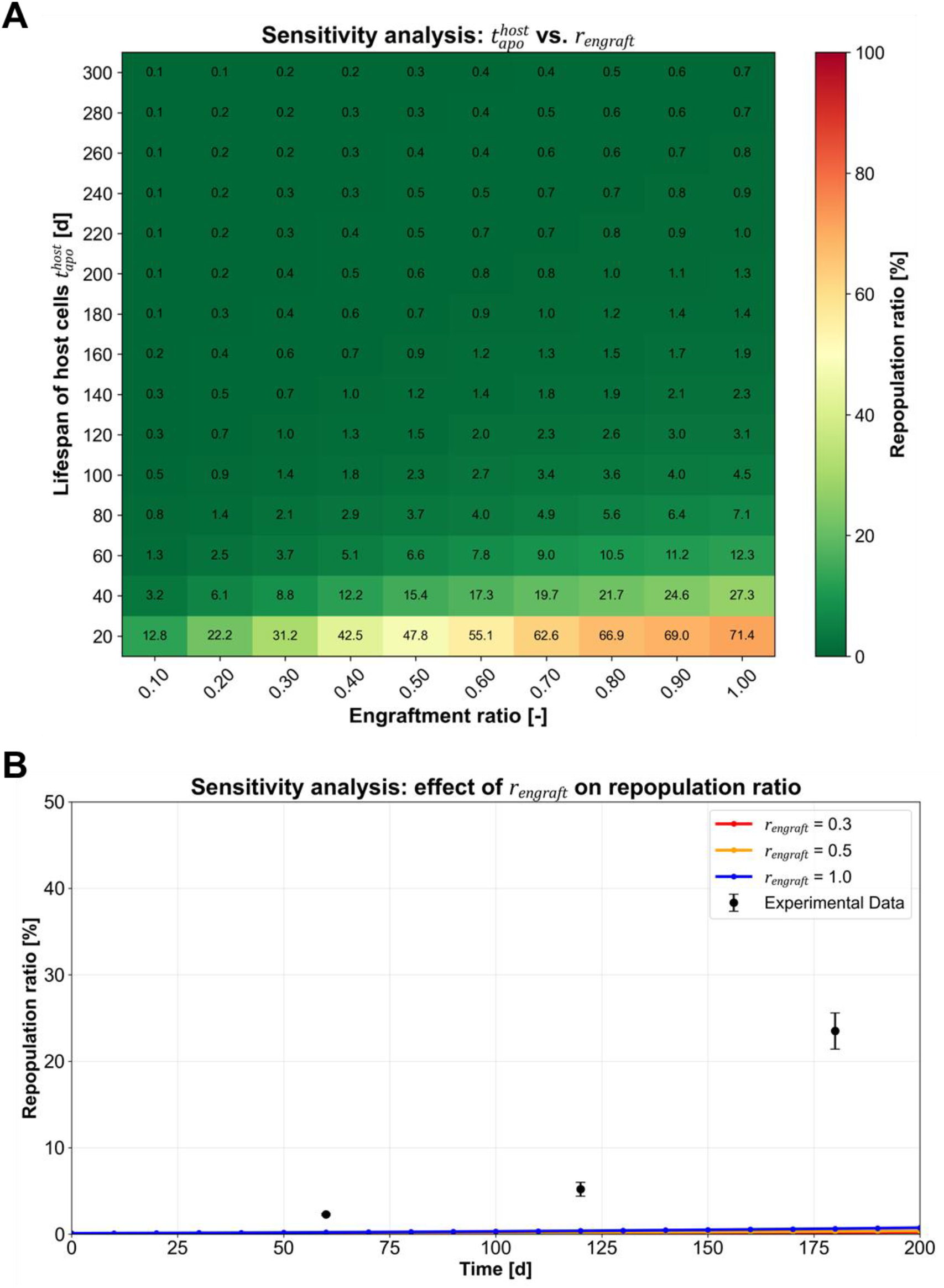
Sensitivity analysis of repopulation efficiency by systemically varying the values of 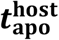 and *r*_*engraft*_. (A) Heatmap of repopulation ratio obtained by varying 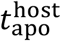 and *r*_*engraft*_. Each grid represents the simulated repopulation ratio. 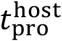 was fixed at 300. (B) Temporal evolution of repopulation ratio for different values of *r*(engraft). 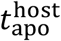 and 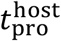 were fixed at 300.

Taken together, these results reject a model in which efficient repopulation is driven solely by passive space competition. Instead, they strongly support the requirement for active cell–cell interactions between transplanted and host hepatocytes.

### 3.2. Estimating the magnitude of cell-cell interaction effects

Motivated by these findings, we introduced a conceptual model incorporating cell–cell interactions, in which host cells are actively eliminated upon contact with transplanted cells. In this framework, host cells acquire an additional probability of apoptosis (*p*_apo_) when adjacent to transplanted cells, independent of their intrinsic lifespan (**Figure 2**).

We then performed sensitivity analysis by varying *p*_interact_, a parameter governing this interaction-dependent apoptosis (see Eq. (5) in Methods). The simulation results indicate that *p*_interact_ = 0.05 is sufficient to recapitulate experimentally observed repopulation dynamics (**Figure 6**).

**Figure 6.**
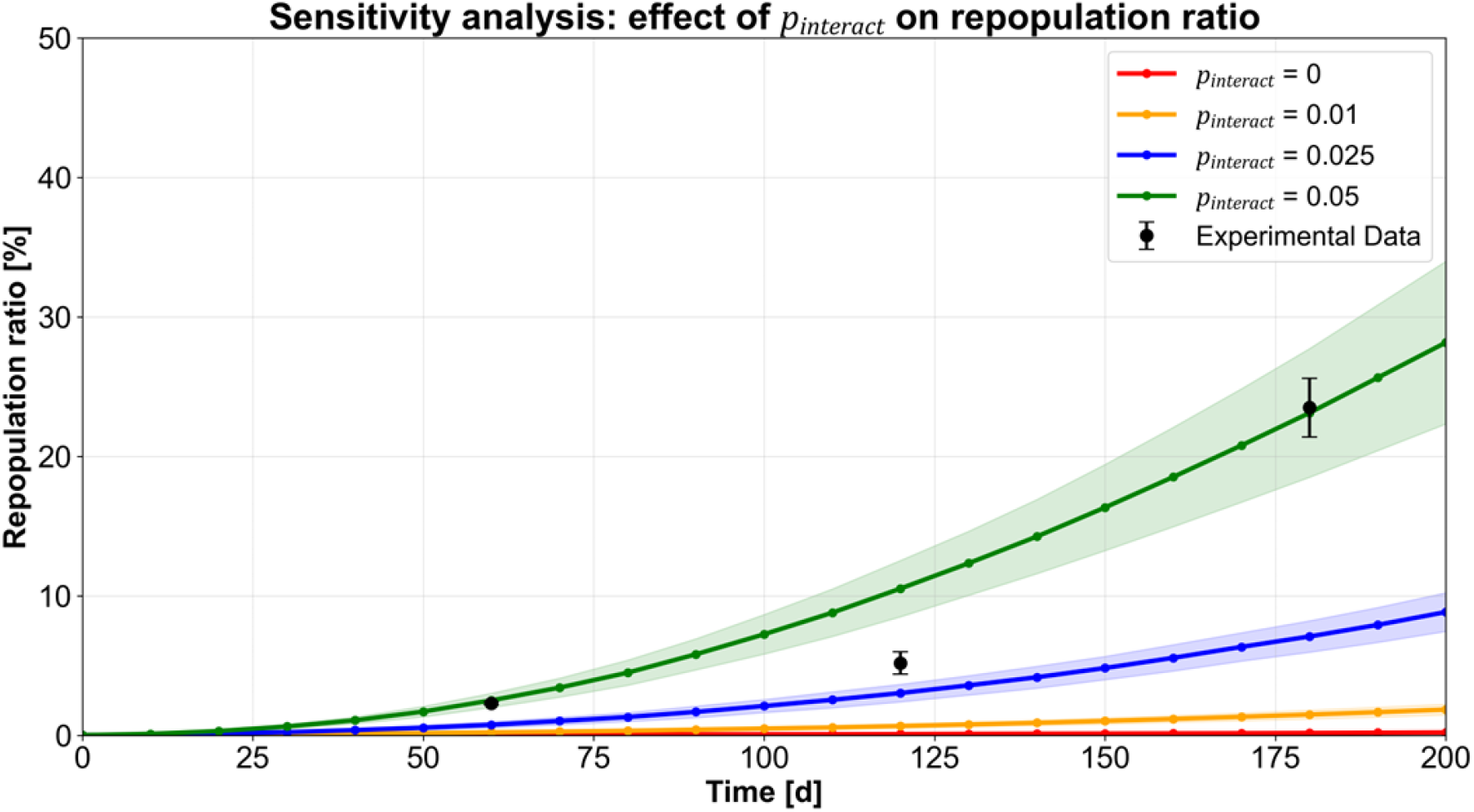
Effect of cell–cell interaction strength on repopulation. Sensitivity analysis of repopulation dynamics with varying *p*_*interact*_, the probability that contact with transplanted cells induces apoptosis in host cells. Temporal evolution of repopulation ratio is shown for different values of *p*_*interact*_ together with experimental data.

Although the molecular mechanisms underlying such interactions remain to be elucidated, this model provides a quantitative framework for estimating the magnitude of cell–cell interaction required to explain repopulation behavior. More broadly, it offers a generalizable approach for dissecting the relative contributions of intrinsic cellular properties and intercellular interactions across diverse biological contexts.

## 4. Discussion

Although earlier studies have suggested a role for cell–cell interactions between host and transplanted hepatocytes in efficient liver repopulation [9,10], their quantitative contribution has remained difficult to assess experimentally. The ABM developed in this study provides a quantitative framework to evaluate whether, and to what extent, such interactions are required to explain observed repopulation dynamics. More broadly, this approach offers a generalizable strategy to disentangle the relative contributions of intrinsic cellular properties and intercellular interactions across diverse biological systems.

This study has several limitations that warrant further investigation. First, key model parameters, including apoptosis rates and proliferative capacity, were estimated from indirect measurements and simplifying assumptions. While such approximations were necessary given the limited availability of experimental data, they may affect the quantitative accuracy of the model. In addition, cell–cell interaction was represented as an abstract, phenomenological term without an explicit molecular basis, which inherently limits its direct biological interpretation. Further elucidation of the underlying molecular mechanisms, together with more precise parameter estimation, will be essential to refine the model and improve its predictive power.

Second, the current analysis focuses on an uninjured host liver, whereas most experimental and clinical settings involve varying degrees of liver injury. In injured livers, host hepatocytes are more prone to apoptosis (i.e., shorter 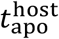), which would increase the contribution of passive selective replacement by transplanted cells. Extending the model to incorporate injury-dependent changes will therefore be necessary to more fully capture the dynamics of hepatocyte repopulation.

Despite these limitations, our results demonstrate that the requirement for cell–cell interaction can be quantitatively evaluated even in the absence of a precise molecular definition. By reducing the problem to a minimal, testable framework, this study provides a conceptual basis for systematically integrating experimental data and progressively incorporating biological complexity. Ultimately, such an approach may help constrain the possible mechanisms underlying cell–cell interaction and guide future experimental investigations.

## Acknowledgements

This research was supported by JSPS KAKENHI Grant Numbers JP24K23176 and JP25K02316, the G-7 Scholarship Foundation Grant and the iPS Academia Japan Grant.

## Author contributions

**Daiki Ikuta**: Conceptualization

**Rei Tamaki**: Conceptualization

**Sota Wada**: Conceptualization

**Kotaro Onishi**: Conceptualization

**Masaki Nishikawa**: Data interpretation

**Yasuyuki Sakai**: Data interpretation

**Takeshi Katsuda**: Conceptualization, supervision

## Notes

### Competing Interest Statement

The authors have declared no competing interest.

